# The psychological arrow of time and the human brain dynamics of event ordering

**DOI:** 10.1101/180711

**Authors:** Baptiste Gauthier, Karin Pestke, Virginie van Wassenhove

## Abstract

When navigating the real-world, the spatiotemporal sequencing of events is intrinsically bound to one’s physical trajectory; when recollecting the past or imagining the future, the temporal and spatial dimension of events can be independently manipulated. Yet, the rules enabling the flexible use of spatial and temporal cognitive maps likely differ in one major way as time is directional (oriented from past-to-future) whereas space is not. Using combined magneto- and electroencephalography, we sought to capture such differences by characterizing time-resolved brain activity while participants mentally ordered memories from different mental perspectives in time (past/future) or space (west/east). We report two major neural dissociations underlying the mental ordering of events in time and in space: first, brain responses evoked by the temporal order and the temporal distance of events-to-self dissociated at early and late latencies, respectively whereas spatial order and distance of events-to-self elicited late brain responses simultaneously. Second, brain responses distinguishing self-position in time and the temporal order of events involved sources in the hippocampal formation; spatial perspective, order and distance did not. These results suggest that the neural dynamics evoked by the temporal ordering of a series of events retrieved from long-term memory, i.e. the psychological time arrow, entails dedicated cognitive processes in the hippocampal formation that are fundamentally distinct from the mapping of spatial location.

## Introduction

When navigating the real-world, the temporal and the spatial order of events being encountered are tightly coupled to one another: the online building of a spatiotemporal context for the encoding of events is naturally tied to the movement of the bodily self moving through its physical trajectory. Brain structures implicated in navigation have been well characterized in the rat and encompass the hippocampal formation [3–5]. Recently, it has been speculated that these regions may also mediate a more abstract from of event mapping in space or time with respect to the self [1]: in the absence of movement of the bodily self, the spatial navigation system has been proposed to support mental operations such as remembering the past or planning the future (i.e. mental time travel) [2] or perhaps, imagining being in a different spatial environment. In other words, the spatial navigation system may be recycled for the spatiotemporal mapping of recollected events, independently of actual movement. Consistent with this working hypothesis, brain regions analogous to the rat’s spatial navigation system have been shown to be active in humans when they navigate in virtual reality in the absence of movement of the bodily self [6–9]. Also consistent with this hypothesis, the functional brain networks mediating spatial navigation in humans show extensive overlap with the memory retrieval system [10]. More specifically, the temporal, spatial and social dimensions of mental events have been shown to engage a common parietal region [11–13] which overlaps with seminal reports of egocentric mapping during actual movement [6].

Altogether, these results suggest that a common operation enabling the setting up of a spatiotemporal context for event retrieval may capitalize on the egocentric remapping of events. Additionally, to make sense in the absence of serial unfolding of events provided by the environment, mental maps need to code for the ordinal attributes of sequences, a property that is essential for the arrow of psychological time [16,17]. While the temporal order of memories may rely on dedicated mechanisms during serial recall [14,18], whether the rules underlying the active ordinal mapping of mental events in their temporal and spatial dimensions share common neural mechanisms is unclear. Previous studies using fMRI found distinct networks subserving temporal order(medial-temporal/prefrontal axis) and spatial location retrieval (Medial temporal/ medial parietal axis) of events from episodic memory [19–21]. However, fMRI studies lacked the temporal resolution to dissociate the sequence of neural events engaged in the cognitive mapping of order.

In the present study (Fig. 1), we combined non-invasive magneto- and electroencephalography (M/EEG) to characterize the precise timing of brain activity indexing the computation of distance and ordinality in the human brain. The structure of the experimental design explicitly separated three cognitive steps necessary to perform the task. First, the participants were asked to mentally imagined themselves away from the ‘here and now’ i.e., in the past or in the future (temporal self-projection, [10]), or to the west or east of their current physical location (spatial self-projection). Second, the participants were informed that an ordinal judgement should be performed in time (TIME task) or in space (SPACE task). Finally, a historical event was presented, and participants judged whether it occurred before/after (TIME) or west/east (SPACE) of the mental self-position in which participants had mentally projected themselves to. For instance, participants were asked to mentally position themselves nine years in the future (or in Cayenne), and judge whether the extinction of elephants happened *before* or will happen *after* (or west/east of) where they mentally were.

**Figure 1.**
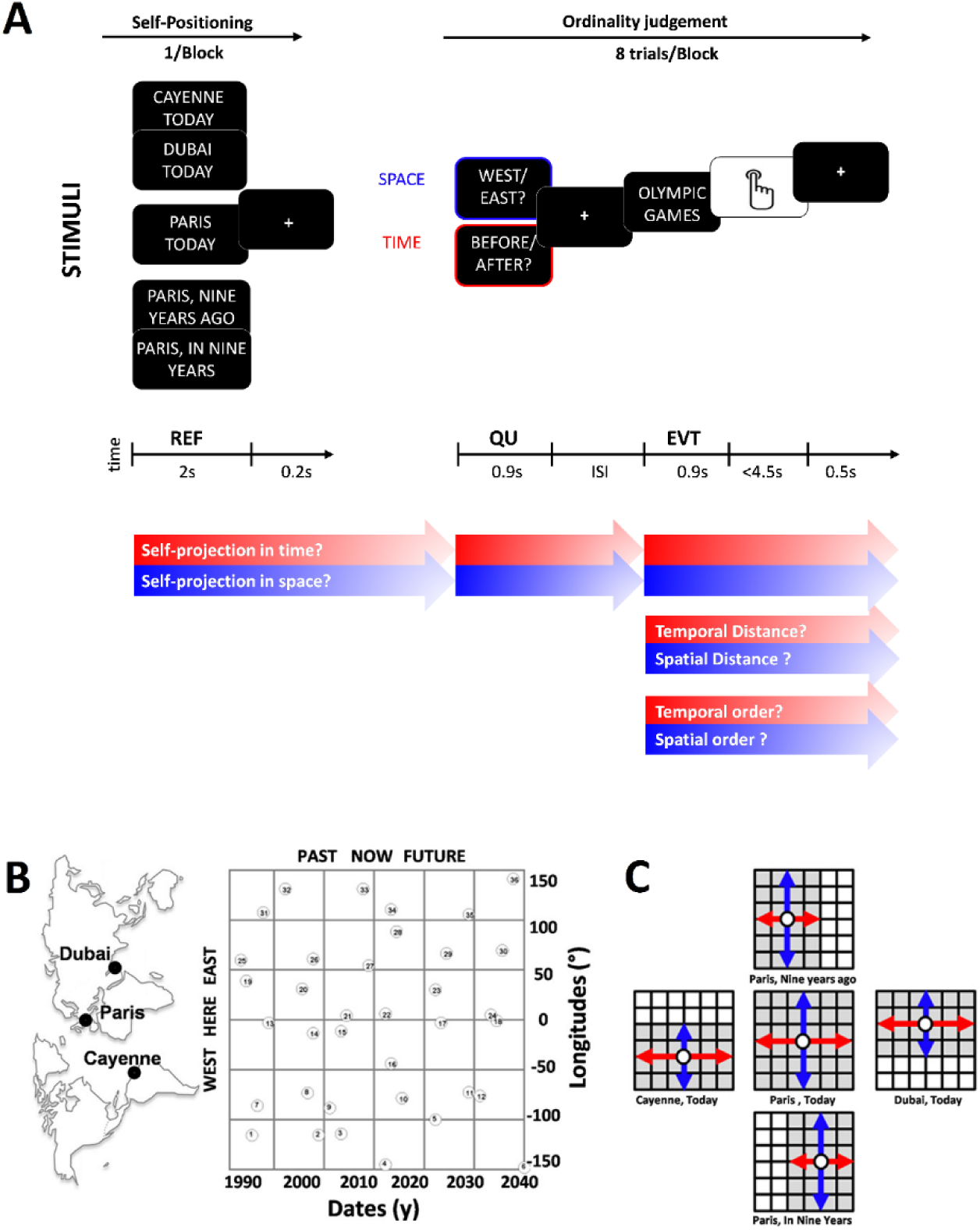
Mental time and space travel with identical historical events. (**A**) The main task consisted in ordering historical events in time (before/after) or space (west/east) with respect to a mental self-position in time and space. The presentation of one of the five possible references (REF) at which participants had to mentally position themselves was followed by a series of eight trials. Each trial started with a question (QU) indicating whether the ordinal judgment would be on the temporal (TIME; red) or on the spatial (SPACE; blue) dimension of events. Any REF could be followed by any dimension (TIME or SPACE) as they were intermixed within a block. Following the QU, one historical event (EVT) selected out of 36 events was presented to the participant. The colored arrows indicate possible moments in brain activity at which markers of self-projection (mental self-position in time or space), distance computation (distance between REF and EVT) and order (position of EVT with respect to REF) may be found. (**B**) Place and date of selected historical events. Events were distributed in a grid-like manner around the possible geographical and temporal locations of the chosen REF. (**C**) To equate the response distribution at chance level, events (shaded grid unit) were distributed so that their relative west/east (blue) and before/after (red) positions were even with respect to each possible REF (open circles). This design ensured a 50% chance level but entailed a restriction of the number of events for positions that were not here (Paris) and now (Today).

Using this experimental design, we observed a sequence of brain responses characterizing the self-projection and the ordering of mental events. Both the latency and the brain regions contributing to these effects were distinctive of the TIME or SPACE tasks. Specifically, temporal ordering of self (self-projection) and events occurred much earlier than spatial positioning, although both involved cortical sources in the hippocampal formation and in medial temporal cortices.

## Results

### Behavioral performance does not inform on ordinality

Behavioral results detailed in fig. S1 were consistent with the hypothesis of common mechanisms for the representation of time and space in the brain. These results were thoroughly discussed in a series of behavioral experiments which uniquely focused on characterizing the cognitive operations implicated in the task ([17]; Experiment 2). Specifically, mapping the temporal or the spatial position of events when mentally imagining oneself away from the ‘here and now’ significantly increased the reaction times (RT) and the error rates (ER). This main effect called the absolute distance effect was specific to the tested dimension so that projecting oneself in time (space) only affected behavioral in the temporal (spatial) task. The second main behavioral effect was the relative distance effect, in which significant decreases of RTs and ERs were found with increasing distance between the mental self-position and the remembered historical event. The absolute and the relative distance effects seminally reported during episodic mental time travel [17] were replicated in this task, and generalized to mental spatial navigation [17,22]. These results supported the hypothesis of comparable self-to-event mapping in time and in space, namely: absolute distance effects were interpreted as indexing self-projection whereas relative distance effects were interpreted as indexing self-to-event distance calculations (Fig. S1A). Crucially, relative distance effects were dimension-specific so that the TIME (SPACE) task was solely affected by the relative temporal (spatial) distances (Fig. S1B). The absence of interferences during the processing of temporal and spatial mapping was shown to engage separate brain regions fMRI [22]. As posited here, the distinct brain regions were likely related to ordinality processing, a major difference between the temporal and spatial dimension. Due to poor temporal resolution of the fMRI technique, no clear effect of ordinality was however found. Furthermore, neither absolute or relative distance effects informed on the direction of mental self-projection, or of the event orientation with respect to the mental self-position: absolute distance effects did not dissociate whether participants performed the task in the past *vs.* future, or in a west *vs.* an east mental self-position; the relative distance effects were also comparable when participants answered ‘before’ or ‘after’ during the TIME task, or ‘west’ or ‘east’ during the SPACE task. Hence, no clear behavioral asymmetries were found as a function of ordinality despite good task performance [17] and no clear neural index of ordinality was yet reported [22]. Here, we thus used time-resolved techniques to tackle what should be a major distinction between time and space, namely, the ordinal arrow of time.

### Self-projection in time precedes self-projection in space

Brain evoked responses following the presentation of the REF were separately averaged as a function of the three possible temporal (PAST, NOW and FUTURE) and spatial (WEST, HERE and EAST) reference points (Fig. 1A, left panel). To investigate the differences evoked by self-projection in time and in space, we performed group-level cluster permutation F-tests separately on the two sets of data.

Self-projection in time showed significant differences at several latencies (Fig. 2A, left panel). An early postero-frontal cluster (MEG-mags: 364 to 1048 ms, p = 0.005), two late clusters (MEG-grads: 564 to 1084 ms, p = 0.003; 1606 to 1829 ms, p = 0.032) and a late frontal cluster (EEG: 1606 to 1829 ms, p = 0.03) dissociated the evoked responses elicited by the three temporal REF. In EEG, the amplitude difference between PAST, NOW and FUTURE was sustained for 3 seconds (fig. 3). These results suggested that self-projection in time occurred as soon as the REF was provided to participants, and may be realized through effortful maintenance over the entire trial. By contrast, no significant F-clusters were found for self-projection in space (Fig. 2A, right panel). To check for potential detectability issues affecting this latter test, we averaged brain activity following a spatial REF (WEST, HERE and EAST) in the clusters detected for self-projection in time, with the ad-hoc hypothesis that self-projection in space and time could entail similar neural substrates. No significant amplitude differences were found using a Bonferroni-corrected pairwise t-test: no evidence for self-projection in space was found following the presentation of the REF.

**Figure 2.**
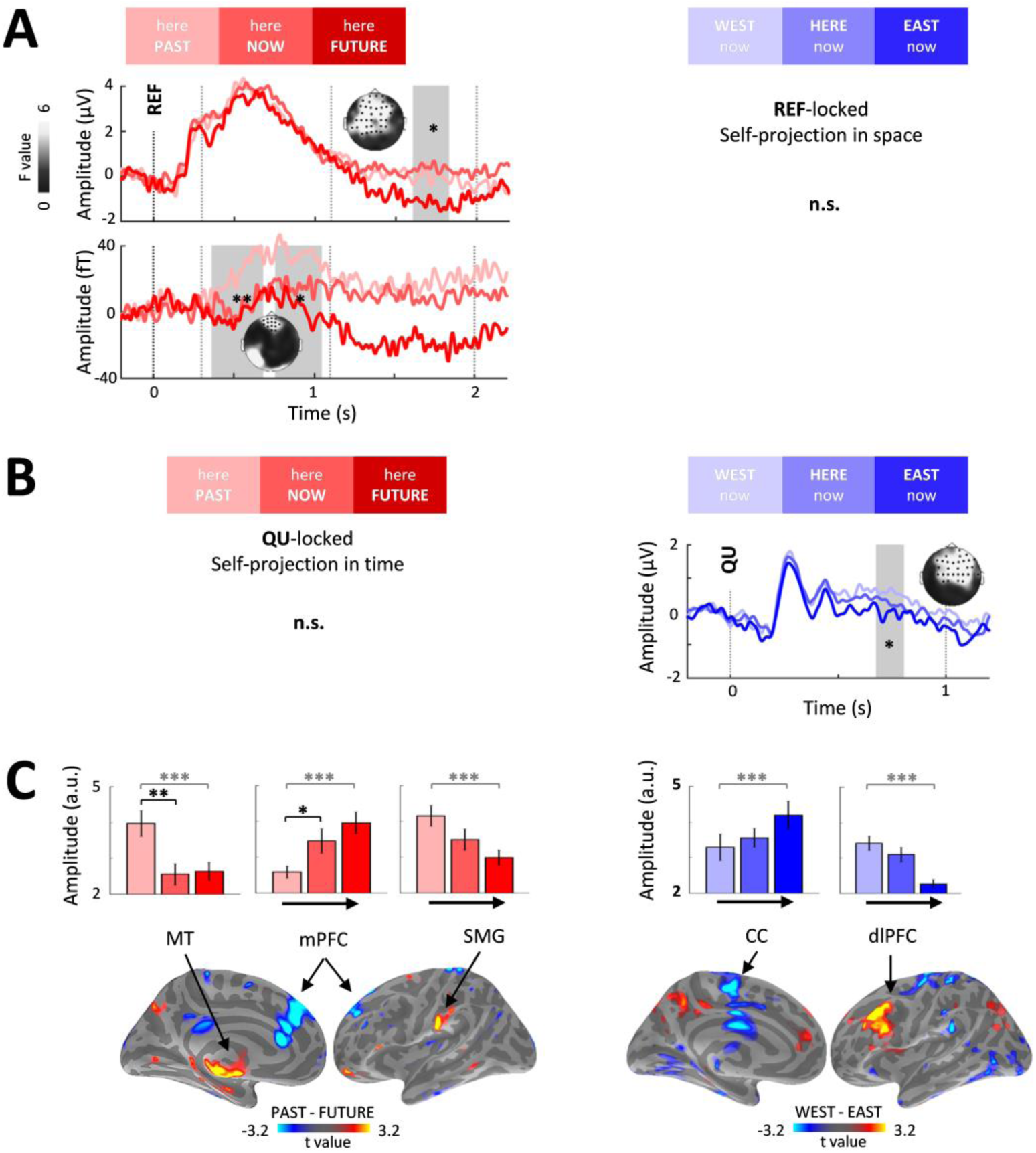
Functional dissociation of self-projection in time (left panels) and in space (right panels). A. Self-projection in time during the presentation of the reference (REF). Left panel: significant differences (shaded areas) recorded with EEG (top) and MEG-mags (bottom) between self-projection in PAST, NOW and FUTURE (light, medium and dark red). Dots on topographical t-maps are significant sensors. Right panel: no significant difference for self-projection in space during REF. **B. Self-projection in space requires task disambiguation (QU).** Left panel: no significant difference between the three temporal REF following the question (QU: before/after?). Right panel: Late brain activity following the question (QU: west/east?) significantly distinguished self-projection in WEST (light blue), HERE (blue) or EAST (dark blue). **C. Separate cortical sources distinguish the direction of self-projection in time and in space**. Left panel: bilateral MT was mostly responsive to self-projection in the PAST (light red); bilateral mPFC and left SMG showed a ranked increase and decrease of activity from PAST to FUTURE references, respectively. Right panel: bilateral CC and dlPFC displayed a ranked increase and decrease of activity from WEST to EAST direction, respectively. Grey stars indicate the significance of whole-brain PAST minus FUTURE (left panel) and WEST minus EAST (right panel) t-tests. Black stars indicate significance of post-hoc t-tests against NOW (left panel) and against HERE (right panel) on clusters first selected by the whole-brain test. Error bars are ± 1 SEM. |t|≥ 3.2 are p ≤ 0.005 for source-level two-sided paired t-tests; clusters > 10 vertices are highlighted*p < 0.05; **p < 0.01.

**Figure 3.**
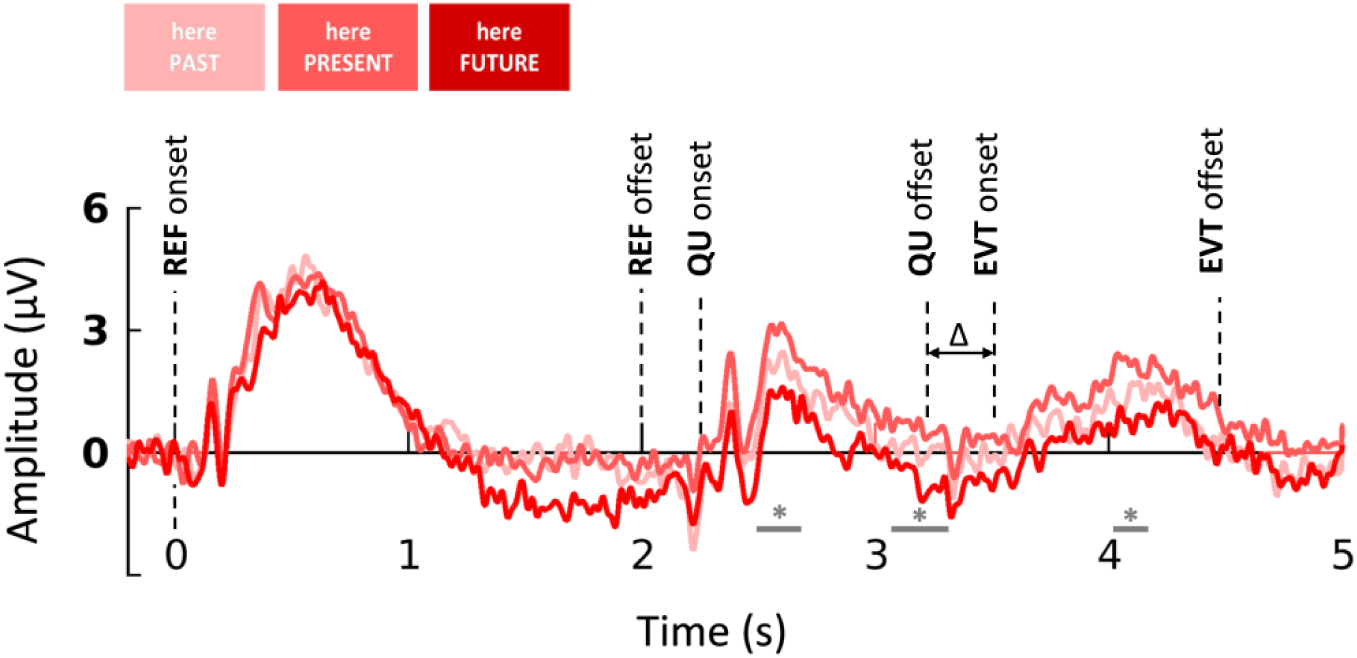
Sustained amplitude differences (EEG) between the PAST, NOW and FUTURE following the presentation of the reference (REF). Differences in evoked responses elicited by the three possible references lasted the entire trial including during the presentation of question (QU) and of the historical event (EVT). The significant clusters indicated by grey bars were found in EEG after the offset of REF at p-values ranging from 0.03 to 0.05 on 2446-2634 ms, 3091-3325 ms and 4008-4138 ms temporal windows This observation suggests that self-projection in time could consist in the maintenance of the mental self-location in time during the whole trial to contextualize the retrieval of the upcoming event. Δ: variable delay between QU offset and EVT onset; * p <= 0.05.

To further investigate brain activity indexing spatial self-projection, brain activity following the presentation of the question (QU-locked brain activity; Fig. 2B) was submitted to a group-level F-test separately for SPACE and TIME. First, QU-locked evoked responses by SPACE trials were sorted as a function of the preceding spatial REF and an F-test revealed a significant EEG cluster (673 to 809 ms, p = 0.009; Fig 2B, right panel). A post-hoc analysis, similar to the one conducted for REF-locked data during self-projection in TIME, was conducted for QU-locked activity during SPACE task, sorted according to the WEST, HERE and EAST reference points. Sensor-level analysis of EEG showed that brain activity in response to the presentation of WEST was significantly higher than that found in response to EAST (t(18) = 4.27, p = 0.0014). The same analysis performed on TIME trials (REF: PAST, NOW and FUTURE) revealed no significant differences (Fig. 2B, left panel). Under the ad-hoc null hypothesis of common brain substrates for self-projection irrespective of the dimension, Bonferroni-corrected pairwise t-tests between QU-locked TIME task evoked responses as a function of PAST, NOW and FUTURE reference points were conducted on the significant sensors found for self-projection in space and showed no significant effects: no evidence for self-projection in time was found following the presentation of the QU.

To clarify the functional role of these differences, we used post-hoc source-reconstruction on the same sorted trials. Based on behavioral absolute distance effects (i.e. increased RT and ER as a function when self-projection), one prediction was that self-projection would yield higher brain activity as compared to no self-projection. Alternatively, self-projection could index the implicit ordering of mental self-positions in time and/or in space as this information was required to accurately perform the explicit order judgments.

### Self-projection is associated with ranked brain activity

Combining the M/EEG contrasts in a common cortical space allowed the estimation of putative brain sources critical for the ranked brain responses observed at the scalp level (Figure 2C). PAST and FUTURE references were contrasted with a vertex-wise paired-t-test averaged over the latency (1606 to 1829 ms) showing the highest difference between the temporal REF across all M/EEG sensors. Three main cortical regions showed significant differences between self-projection in PAST and FUTURE (Fig 2C; Fig S2 and S3; MNI coordinates in table S1): bilateral medial temporal (MT), bilateral medial prefrontal cortex (mPFC), and the left supramarginal gyrus (SMG). Using Bonferroni-corrected pairwise paired t-test contrasting brain activity between PAST and FUTURE *vs*. NOW revealed significantly higher activity for PAST than for NOW (Fig. 2C, left panel) in MT (t(18) = 3.86, p < 0.005) and mPFC (t(18) = 2.84, p = 0.032) whereas SMG showed a decrease of activity from PAST, to NOW to FUTURE. Inspection of the temporal course of the source estimates confirmed that the significant ranked effect of self-position in time emerged 1 s following the presentation of the REF (Fig. S3).

The same approach was taken for QU-locked source estimations in SPACE (Fig. S4 and S5). Source estimates for self-projection in space were investigated with the contrast WEST *vs*. EAST that most contributed to the significance of F-test in sensors (Fig. S4). Two main regions showed significant effects of the spatial REF on brain activity following the disambiguation of the dimension to be judged (Fig. 2C right panel; MNI coordinates in Table S1): bilateral dorsolateral prefrontal cortices (dlPFC) and cingulate cortices (CC). Following the presentation of QU, the dlPFC showed ranked activity according to the direction of the REF from WEST to EAST whereas CC showed ranked activity going from WEST to EAST (Fig. 2C). Both regions showed an ordering of the source estimate amplitude as a function of the imagined self-position in space. Bonferroni-corrected pairwise t-test did not reveal significant differences between WEST and HERE or between HERE and EAST. The ordinality effect of self-position in space emerged as early as 300 ms following the presentation of the QU and remained sustained throughout the presentation of the question (Fig. S4). We then questioned whether brain responses indexing explicit ordinal judgments would be elicited by the presentation of the event (EVT-locked activity).

### Sequential processing of temporal order and distance; parallel processing of spatial order and distance

EVT-locked responses were sorted according to two metrics: an ordinality metric capturing the signed distance between the imagined self-position (REF) and the EVT (e.g., 5 years before the REF corresponded to a signed distance of −5 in the TIME task), and an distance metric, in which the ordinal relation between REF and EVT was disregarded by taking the absolute value (e.g., 5 years before or after the REF was assigned a value of 5). Separate subject-wise regressions of EVT-locked brain activity were performed with signed and absolute distance, separately. Group-level cluster two-sided permutation t-tests were then performed on the resulting single-participant regression betas. These analyses were performed separately for the TIME and SPACE tasks with their respective signed and absolute distance metrics.

In TIME task, early clusters significantly correlated with the signed temporal distance (MEG-mag: 372 to 520ms, p = 0.015; MEG-grad: from 444 ms to 544 ms and from 592 ms to 724 ms, p = 0.002). Over the common significant time window (444 to 520 ms), the further an event was in the past, the more negative the amplitude of the evoked response irrespective of the mental self-reference; conversely, the more distant an event was in the future, the more positive the evoked response. The amplitude of the evoked responses was thus indicative of the ordinal position of the EVT in time with respect to the mental self-position (Fig. 4A, top). This pattern of brain activity captured ordinality processing, which was not observed in behavioral results (fig. S1). Conversely, consistent with the symmetry of RTs an ERs in behavior, clusters significantly correlating with the absolute value of temporal distance were also found at late latencies (MEG-mag: 828 ms to 1000 ms, p = 0.006; MEG-grad: 440 ms to 500 ms, p = 0.001 and 840 ms to 1000 ms, p = 0.008), whose amplitude showed a pattern indexing the temporal distance effect with respect to the REF: the further the event was from the REF, the more positive the amplitude of the evoked response (Fig. 4A, bottom).

**Figure 4.**
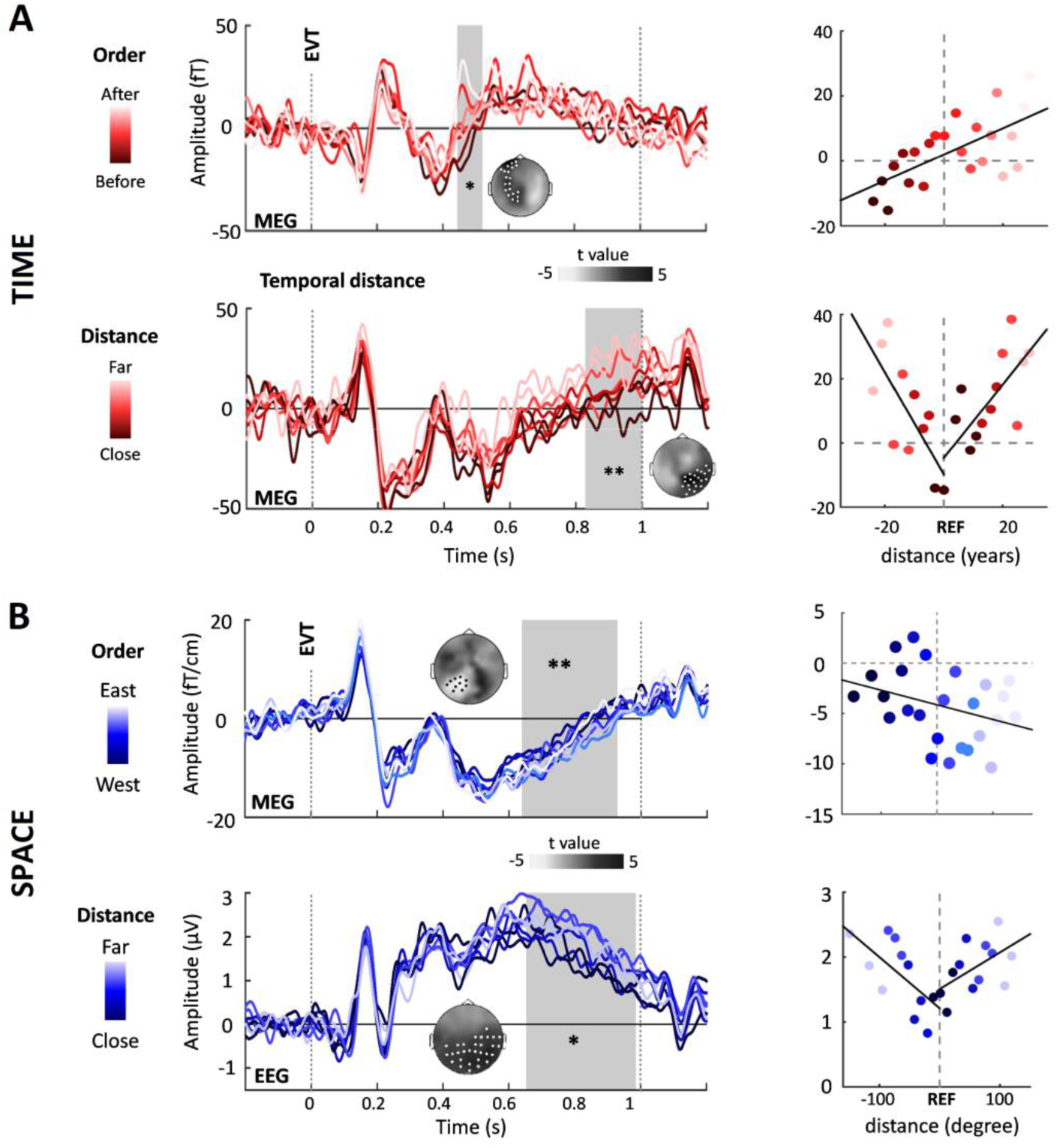
Order and distance are sequentially processed in TIME (red) and simultaneously in SPACE (blue). Zero is the presentation of the EVT; EVT-locked responses were sorted as a function of 8-binned metrics. Significant cluster topographies are displayed alongside with a gray area indicating their timing**. (A) TIME task sequential effects.** Top left: temporal order effect timecourse. MEG-mags activity at 444-520 ms correlates with the ordinal position of the historical EVT with respect to the REF. Top Right: fined-grained effect. Mean cluster activity per single ordered distance value: the more positive the activity, the later (lighter red) the events; the more negative the activity, the earlier (darker red) the EVT. Bottom left: temporal distance effect timecourse. MEG-mags activity at latencies 828-1000 ms correlated with the absolute temporal distance separating the EVT from the REF. Bottom right: fined-grained effect. Mean cluster activity per single absolute distance value at the latency of interest. The more positive, the further (lighter red) the event was from the REF. **(B) SPACE task simultaneous effects.** Top left: spatial order effect timecourse. MEG-grads activity over 640-928 ms correlate with the ordinal position of historical events with respect to the spatial REF. Top Right: fine-grained effects. Mean cluster activity per single ordered distance value at the latency of interest: the more negative the activity, the more eastern (light blue) the EVT. Bottom left: spatial distance effect timecourse. EEG activity over 653-985 ms correlated with absolute spatial distance to the REF. Bottom right: fine-grained effects. Mean cluster activity per single absolute distance value at the latency of interest. The more positive the activity, the further (lighter blue) the events were from the REF. * p < 0.05; ** p < 0.01.

The same regression analyses carried out for spatial metrics in SPACE task revealed significant late clusters correlating with the signed spatial distances (MEG-grad: 640 ms to 928 ms, p = 0.019) so that the response amplitude indexed the west/east spatial location of the EVT with respect to the mental self-position. The further the event was in the relative west, the less negative the amplitude of the evoked response; the further the event was in the relative east, the more negative the amplitude of the evoked response (Fig. 4B, top). Similar to temporal effects, significant clusters correlating with the absolute value of spatial distance were found (EEG: 653 ms to 985 ms, p = 0.002) so that the amplitude of brain responses indexed spatial distances with respect to the REF: the further the event was from the reference, the more positive the evoked response (Fig. 4B, bottom).

### Distinct source estimates for order and distance computations

EVT-locked activity over the latency of interest for sensor-level effect of temporal order (444 to 520 ms; Fig. 4A, top) showed a significant linear regression of amplitude with the signed distance in bilateral MT (Figure 5A, left; Table S1). Additionally, source estimations of temporal distance cluster showed a significant linear regression of source amplitude with the binned absolute value of temporal distance in left retrosplenial cortex (RSC), left inferior frontal gyrus (IFG) and left dlPFC (Fig. 5A, right; Table S1). Conversely, for spatial order effects, the amplitude of source estimates showed a significant linear regression with binned spatial order in left dorsomedial prefrontal cortex (dmPFC), intraparietal sulcus/superior parietal lobule (IPS/SPL) and anterior insula (aINS) (Fig. 5B, left; Table S1). Additionally, source-level activity showed a significant linear regression between the amplitude of the sources estimates in left precuneus (PCUN) and posterior parahippocampus (PHC) with binned absolute value of spatial distance (Fig. 5B, right; Table s1).

**Figure 5.**
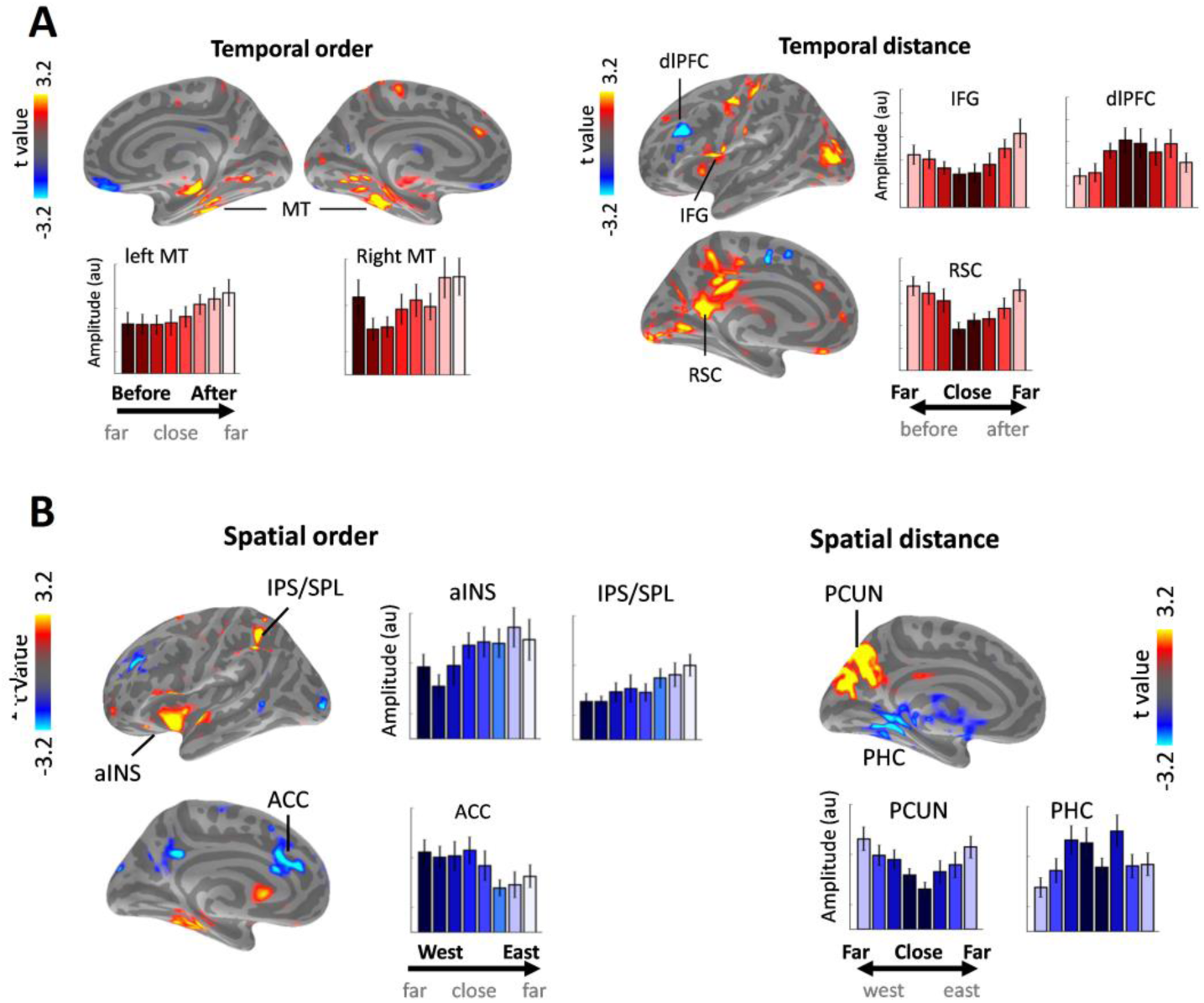
Distinct networks generate distance and order effects in TIME and SPACE. (A) Time task sources. The source estimations of the temporal order effect (left) showed a monotonic increase of activity in bilateral MT with the 8-binned signed distances. By contrast, source estimations of the temporal distance effect (right) revealed a monotonic increase in RSC and IFG and a monotonic decrease in dlPFC with the 8-binned temporal distance. **(B) Space task sources.** Source estimates of the ordinality effect showed a monotonic increase of activity in left IPS/SPL and aINS and a monotonic decrease in ACC with the 8-binned signed spatial distance. By contrast, source estimates of the distance effect showed a monotonic increase in PCUN and a monotonic decrease in PHC with 8-binned absolute spatial distance. Error bars correspond to ± 1 SEM. |t|≥ 3.2 correspond to p ≤ 0.005 for source-level two-sided paired t-tests, cluster > 10 vertices are highlighted.

## Discussion

Our results suggest a fundamental distinction between the conscious mapping of events in time and in space, highlighting the particular status of ordinality for the psychological arrow of time. In the task used here, a fundamental computational goal to correctly perform ordinal judgements was to represent an imagined self position in time [10,16,17] or in space. We thus propose that this mental self-position may act as the anchor (or context) for the computation of ordinal position [17] effectively yielding self-to-event computations of order and distance.

### Early encoding of self-positions in time in a self-processing network

Long before event retrieval (EVT), and immediately following the presentation of a temporal REF, the cortical source estimates contributing to self-projection in TIME (medial temporal, medial prefrontal and right supramarginal cortices) were consistent with regions found with fMRI during mental time travel [23–25] and temporal order tasks [26,27]. We found that the activity estimated in the medial temporal cortex was mostly driven by the past, consistent with previous fMRI work [23,28], suggesting that self-projection in the past was accompanied by the recollection of past events. By contrast, the engagement of frontal regions has previously been reported to be more important during the imagination of the future than the remembrance of the past [25,29,30]. In our study, using PAST, NOW and FUTURE condition enabled to parametrically investigate the representation of the time arrow as a function of self-projection: activity estimated originating in frontal cortex was found to index the temporal succession of past, present and future. To the best of our knowledge, this is the first time-resolved study showing that that temporal self-position evoked a gradual activity in frontal cortices during past, present and future self-projection. Incidentally, we also found that evoked responses source-estimated to the right inferior parietal lobule (SMG) were larger in the NOW compared to the PAST and FUTURE conditions, which is consistent with previous observations of higher temporo-parietal BOLD activity related to present compared to self-projection in the past and the future [22]. Both IPL and medial frontal cortices have been generally involved in self-processing [32,33] and in self-processing during episodic memory [34,35], suggesting that the representation of self-position in time could be a fundamental aspect of self-processing and possibly a basis for the experienced continuity of identity over time.

The sequences of brain responses over time suggest that participants actively build a temporal referential, in medial frontal regions, to locate retrieved events [16,17]. In the mental time travel task used here, historical events appeared to be automatically contextualized by previous changes in mental self-position, but imputed an ordinal position with respect to this mental reference, at a later decisional stage. The immediate ranking of the amplitude of brain evoked responses in mPFC and SMG during the presentation of the REF (Fig. 2) suggest that the directionality of time, i.e. the succession of past, present and future, may be set very early on and this contextualization remained active throughout the entire trial (Fig. 3). By contrast, self-projection in SPACE occurred much later, following the presentation of the question, which disambiguated the temporal *vs.* spatial dimension of the task. This observation supports the *active suppression hypothesis* positing the competition between the temporal and the spatial mapping of events (Cf. Supp. Mat.; [17]) and would with consistent with the early latencies of the self-projection in time effect, with a prioritization of temporal over spatial processing. Regions contributing to self-projection in space included the core spatial attention network [36,37] indicating the possible use of a visuospatial strategy to self-project in space prior to event retrieval.

Overall, in both TIME and SPACE, and without using particularly tailored test to detect the ranking of brain response amplitude, we observed ranked brain responses indexing the mental self-position with respect to the ‘here and now’. These results indicate that self-projection may generally involve an implicit ranking for where the self is located in time and space. Mental time appears special in that it is prioritized over mental space and this likely relates to more conscious aspects of self-representation.

### Sequentiality of temporal metrics processing as compared to space

Following the presentation of the event, the early latency of the temporal ordinality effect (Fig. 4A, top) and its corresponding cortical source estimates (Fig. 5A, left) in medial temporal regions were compatible with mechanisms involved in temporal sequencing [14,18,38] and temporal distance [39] computations during the retrieval of events. Furthermore, the explicit and egocentric ordinal classification, but not the distance effects proper, involved the hippocampal formation (Fig. 5A, left): ordinality was indexed earlier in time than the distance effects, the latter likely reflecting a decisional stage (Fig. 4A; fig. S6). Our time-resolved neuroimaging results suggest that the cognitive processing of ordinality in mental time travel unfolds in three sequential steps: ordinal contextualization, egocentric retrieval, and computation of egocentric ordinal relations between context and event. One possible alternative would be that the correlation between ordinal distance and MT activity index a memory load effect [40], indicating that our task was equivalent to a memory search in a list of events and consequently, the ordinal position of event would be reducible to a serial position of an item in a list. However, we content that this is highly unlikely considering our pattern of results. First, no primacy or recency effects (typical effects of seriality in memory of lists) were observed in our task. Second, all our behavioral and M/EEG effects are *relative* to the self-position: whether the reference was nine years in the past, the present time or nine years in the future, the behavioral patterns were the same [17]. In other words, our results suggest that the serial position in time is here egocentric, centered on the imagined self-position and not absolute as would be expected in a list search. Additionally, we found absolute temporal distance correlates in RSC and frontal cortices, consistently with localizations reported in previous fMRI work [41]. Because RSC and frontal cortices are functionally coupled to MTL during self-relevant memory [42,43], the absolute temporal distance effects observed here could relate to the local self-relevance of events (the closer, the more relevant).

By contrast, self-projection in space displayed a ranked activity mirroring the spatial alignment of possible references (Cayenne, Paris and Dubaï) only after the presentation of the question, when participants expected a subsequent spatial ordinal judgment. The cortical source estimates for this effect, combined with explicit knowledge of the spatial dimension for the task, suggested that participants used a cognitive strategy akin to top-down visuospatial attention oriented towards internal representations [37,44]. Additionally, the neural correlates of spatial ordinal processing were concurrent with the spatial distance effect, indicating a parallel processing of distinct spatial metrics. Superior parietal regions are typically observed during spatial egocentric processing of spatial information for navigation [45–48], consistent with the idea of a mental transformation of event position in a new spatial reference frame [49]. The pattern found for spatial distances in our study fits well with the local/global spatial framework developed for spatial navigation [50], and is overall consistent with the idea of the recycling for mental travels of computations used in real-world navigation. By contrast, temporal order processing appeared to engage dimension-specific cognitive processes.

## Conclusion

On the basis of these empirical findings, we propose that mental time travel relies on a specific sequence of neural computations emphasizing ordinality between the cognitive self and remembered events, as compared to a controlled matched spatial task *a priori* implicating very similar algorithmic steps. The medial temporal - prefrontal axis provides a reference frame for the self in time [22,51], allowing subsequent fast processing of self-related temporal order in hippocampus [1]. In sum, we hypothesized a distinct series of operations engaged in mental travel tasks and showed possible neural activity dedicated to the building of a conscious psychological arrow of time.

## Material and Methods

**Stimuli and procedure.** At least 48 h before the main neuroimaging experiment, participants were provided with a list of historical events, which each of them provided with a short historical context, its date, and its location on a world map centered on Paris (https://www.google.ca/ maps, June 1, 2013). During the M/EEG experiment, stimuli consisted in viewing white words projected on a screen placed 90 cm away from participants seated in a magnetic-shielded room, under the Neuromag Elekta MEG system (LTD, Helsinki). Participants were asked to imagine or mentally project themselves to a reference point in time or space. Five possible references (REF) were tested: ‘9 years ago, Paris’ (PAST), ‘Today, Paris’ (NOW, HERE), ‘in 9 years, Paris’ (FUTURE), ‘Today, Cayenne’ (WEST) and ‘Today, Dubaï’ (EAST). The NOW and HERE were control references as no self-projection were expected in the temporal or the spatial dimension. Self-projections in time (PAST, FUTURE) did not entail self-projection in space (both were in Paris); self-projections in space (WEST, EAST) did not entail self-projection in time (both were Today). Hence, the temporal and spatial dimensions of self-projection were tested separately. In a given block (8 consecutive trials), the REF was displayed on the screen for 2 seconds followed by a fixed delay of 200 ms. One REF was shown every eight trials (Fig. 1A). Following the presentation of the REF on the initial trial of a block, and at the beginning of every subsequent trials, participants were prompted with a question instructing them to perform an ordinality task either in the temporal or in the spatial dimension. On a given trial, the question (QU) prompted participants to choose in a two-alternative-forced choice whether the historical event was before or after (‘avant/après’; TIME) or west or east (‘ouest/est’; SPACE) of the mental self-position (Fig. 1A). The QU lasted 900 ms on the screen, followed by a short variable ISI ranging from 220 to 350 ms, and by the historical event (EVT) displayed for 900 ms on the screen. Hence, the historical event on which the ordinality judgement had to be made was presented after the QU. Following the presentation of the EVT and a blank screen of 500 ms, participants were given up to 4.5 s to provide their answer before the next trial automatically started. The experimental instructions emphasized accuracy and speed equally.

**Simultaneous MEG and EEG recordings.** Simultaneous magnetoencephalography (MEG) and electroencephalography (EEG) were recorded to improve both the spatial coverage and the sensitivity to a diversity of possible cortical and subcortical generators. Continuous MEG and EEG were simultaneously collected using a whole-head MEG system with 102 magnetometers (MEG-mags) and 204 planar gradiometers (MEG-grads) (Elekta Neuromag Vector View 306 MEG system) and the built-in EEG system (60 channels) at a sampling rate of 1000 Hz and bandpass filtered between 0.03 Hz and 330 Hz. Participants were seated in an upright position. The head positions of participants were measured before each block with four head position coils (HPIs) placed over the frontal and mastoid areas. Three fiducial points (nasion, left and right pre-auricular areas; Polhemus Isotrak system) and EEG electrodes positions were used during the digitization for later coregistration with anatomical MRI. Electro-occulograms (EOG, horizontal and vertical eye movements) and electrocardiogram (ECG) were simultaneously recorded with M/EEG.

**Event-related potentials/fields in sensor space.** All analysis were done in compliance with current guidelines in the field [52] M/EEG signals were downsampled to 256 Hz and low-passed filtered at 35 Hz prior to epoching. M/EEG trials were epoched per condition of interest and locked on the onset of the word (REF, QU or EVT) associated with the experimental condition of interest. All epochs were baseline corrected using the 200 ms prestimulus period. In the main text, we refer to evoked activity indistinctly of the method of recording (MEG-mag, MEG-grad and EEG are specified in the statistical results), and use REF-locked for averages with the reference as the stimulus at zero ms, QU-locked for averages locked on the question and EVT-locked for averages locked to the event.

The significance of group-level contrasts was assessed using non-parametric cluster permutation procedure using Monte-Carlo procedure (Fieldtrip toolbox [53]). 1000 permutations were used to determine significance; the seed was fixed for reproducibility. Only spatio-temporal clusters with corrected p-values ≤ 0.05 are reported for F-tests, and ≤ 0.025 for two-sided t-tests. Contrasts performed on REF-locked brain activity were split in two time windows ranging from 300 to 1100 ms and from 1000 to 2000 ms in order to equate the number of data points in the different time windows of analysis as spatiotemporal clustering technique is sensitive to this parameter[53]. To investigate the effect of self-projection, F-tests were separately performed on REF-locked data as a function of the 3 possible references in time (PAST, NOW and FUTURE) or in space (WEST, HERE and EAST). Then, F-tests were separately performed on QU- and EVT-locked data as a function of the 3 possible references in time (PAST, NOW and FUTURE) in TIME task, and the 3 possible references in space (WEST, HERE and EAST) in SPACE task.

To investigate the effect of ordinality on one hand and distance on the other hand, to two distance metrics were used. First, an ordinality metric captured the signed distance between the imagined self-position (REF) and the EVT (e.g., 5 years before the REF corresponded to a signed distance of −5 in the TIME task). Second, a distance metric captured the absolute value of distance, in which the ordinal relation between REF and EVT was disregarded (e.g., 5 years before or after the REF was assigned a value of 5). EVT-locked data were first averaged across the 24 distance bins. The temporal distances were used for TIME and the spatial distances for SPACE. A subject-wise linear regression between the amplitude of the evoked responses and the (temporal or spatial) distance was computed for each sensor and each time point. A group-level two-tailed one-sample t-test was then performed on the regression betas. The same procedure was applied with the absolute value of the distance.

**Combined M/EEG source reconstruction.** The co-registration of MEG and EEG data with an individual’s anatomical MRI was carried out by realigning the digitized fiducial points with the EEG electrodes. We used a two-step procedure to insure reliable co-registration between MRI and MEG coordinates: using MRILAB (Neuromag-Elekta LTD, Helsinki), fiducials were aligned manually with the multimodal markers visible on the MRI slice; an iterative procedure was then used to realign all digitized points with the scalp tessellation using the mne_analyze tools within the MNE suite. Individual forward solutions for all source reconstructions located on the cortical sheet were next computed using a 3-layer boundary element model [54,55] constrained by the individual anatomical MRI. Cortical surfaces were extracted with FreeSurfer and decimated to about 10240 vertices per hemisphere. The noise covariance matrix for each individual was estimated on baseline segments ranging from 200 to 0 ms before stimulus onsets. The inverse computation was done using a loose orientation constraint (loose = 0.2, depth = 0.8) (Lin et al., 2006). The reconstructed current orientations were pooled by taking the norm, resulting in manipulating only positive values. For each condition of interest, a standard smoothing was applied prior to group-level statistics computation (smooth parameter = 20).

For source space statistics analysis, epochs were imported in mne-python [56]. For each contrast, the number of epochs per condition was equalized prior to source reconstruction to avoid source amplitude bias caused by different signal-to-noise ratios, using the mne.equalize.epochs_count method in mne_python library. This method seeks to reduce the impact of time-varying noise by minimizing the differences in the times of the events in the two or more sets of epochs that have to be equalized. Source-level contrasts for self-projection in time and in space were chosen to be (PAST-FUTURE) and (WEST-EAST), respectively. These were chosen according to the results of the F-tests between PAST, NOW and FUTURE and WEST, HERE and EAST (see Main Text and supplementary fig. S2 and fig. S4). Source-level distance contrasts were obtained as follows: epochs were averaged across 8 distance bins (to improve the SNR compared to the 24 bins in sensor space) and subject-wise linear regressions were performed between source estimate amplitudes and binned distance. For every contrast, a group-level two-sided one-sample paired t-test was performed for each vertex on the differences between conditions, or on the regression betas. A source-space contrast cluster was considered significant if more than 12 adjacent vertices survived a p ≤ 0.005 uncorrected threshold in order to balance the stringency and the sensitivity of source-space analyses.

## Author contributions

BG and VvW designed the experiment, BG and KP acquired the data, BG and VvW analyzed the data and BG and VvW wrote the paper.

## Acknowledgments

This work was supported by an ERC-YStG-263584 and an ANR10JCJC-1904 to V.vW. We thank the members of UNIACT and the medical staff at NeuroSpin for their help in recruiting and scheduling participants. We thank members of UNICOG for fruitful discussions. Preliminary data were presented at SFN (2014, Washington DC), OHBM (2015, Geneva) and CNS (2016, New York).

